# Microarray profiling of hypothalamic gene expression changes in Huntington’s disease mouse models

**DOI:** 10.1101/2022.03.15.484411

**Authors:** Elna Dickson, Amoolya Sai Dwijesha, Natalie Andersson, Sofia Lundh, Maria Björkqvist, Åsa Petersén, Rana Soylu-Kucharz

**Affiliations:** Biomarkers in Brain Disease, Department of Experimental Medical Science, Lund University, Sweden; Translational Neuroendocrine Research Unit, Department of Experimental Medical Science, Lund University, Sweden; Pathways of cancer cell evolution, Division of Clinical Genetics, Department of Laboratory Medicine, Lund University, Sweden; Department of Pathology and Imaging, Global Drug Discovery, Novo Nordisk A/S, Denmark

**Keywords:** Huntington’s disease, neuroendocrine, hypothalamus, microarray, HD mouse models, LIMMA-GSEA, pathway enrichment analysis, qRT-PCR, overexpression, huntingtin

## Abstract

Structural changes and neuropathology in the hypothalamus have been suggested to contribute to the non-motor manifestations of Huntington’s disease (HD), a neurodegenerative disorder caused by an expanded CAG repeat in the huntingtin (HTT) gene. In the present study, we investigated whether transcriptional changes would be part of hypothalamic pathology induced by the disease-causing huntingtin (HTT) protein. We performed microarray analysis using the Affymetrix platform on total hypothalamic RNA isolated from two HD mouse models and their littermate controls; BACHD mice with ubiquitous expression of full-length mutant HTT (mHTT) and wild-type mice with targeted hypothalamic overexpression of either wild-type HTT (wtHTT) or mHTT fragments. To analyze microarray datasets (34760 variables) and obtain functional implications of differential expression patterns, we used Linear Models for Microarray Data (limma) followed by Gene Set Enrichment Analysis (GSEA) using ClusterProfiler. Limma identified 735 and 721 significantly differentially expressed genes (adjusted p < 0.05) in hypothalamus of AAV datasets wtHTT vs control and mHTT vs control. In contrast, for BACHD datasets and the AAV mHTT vs. wtHTT dataset, none of the genes were differentially expressed (adjusted p-value > 0.05 for all probe IDs). In AAV groups, from the combined limma with GSEA using ClusterProfiler, we found both shared and unique gene sets and pathways for mice with wtHTT overexpression compared to mice with mHTT overexpression. mHTT caused widespread suppression of neuroendocrine networks, as evident by GSEA enrichment of GO-terms related to neurons and/or specific neuroendocrine populations. Using qRT-PCR, we confirmed that mHTT overexpression caused significant downregulation of key enzymes involved in neuropeptide synthesis, including histidine and dopa decarboxylases, compared to wtHTT overexpression. Multiple biosynthetic pathways such as sterol synthesis were among the top shared processes, where both unique and shared genes constituted leading-edge subsets. In conclusion, mice with targeted overexpression of HTT (wtHTT or mHTT) in the hypothalamus show dysregulation of pathways, of which there are subsets of shared pathways and pathways unique to either wtHTT or mHTT overexpression.

## Background

Huntington’s disease (HD) is a fatal neurodegenerative disorder caused by a mutation in the huntingtin (*HTT*) gene, which results in a CAG repeat expansion in exon 1 of *HTT* [1]. The expanded repeat leads to the formation of a mutant protein (mHTT) with an abnormally long polyglutamine (polyQ) stretch, resulting in protein misfolding and aggregation of the mutant protein in neurons [1, 2]. The length of CAG repeats is inversely correlated to the age of onset, with 40 or more CAG repeats resulting in full-penetrance and rapid-onset of HD, while 60 or more repeats lead to juvenile HD [3, 4]. Currently, there is no cure for HD, and a broad range of clinical trials aims to directly silence HTT, including interventions that either silence both alleles (normal and mutant) or selective targeting of the mutant protein [5, 6]. This, however, stresses the need for further understanding of the function of normal HTT in cells. Several *in vitro* and *in vivo* studies showed that normal HTT is involved in numerous cellular processes, including cell maturation, vesicle trafficking, synaptic transmission, and neuroprotection (reviewed in [7-10]). Additionally, further knowledge about how gains- or losses-of-dysfunctional mHTT affects the various tissues and cell types throughout the body is needed.

The hypothalamus is the central brain region for the regulation of energy metabolism and plays the main role in the central-peripheral regulatory network that maintains body homeostasis [11-14]. Imaging and post-mortem studies in clinical HD and HD animal models have shown pathological changes such as atrophy, reduced grey matter content, and loss of neuropeptides in the hypothalamus, some that can be detected even before the onset of motor features [15-20]. Metabolic alterations and other non-motor symptoms with a hypothalamic link are present throughout all stages of HD [21-24] and a higher baseline body mass index (BMI) is associated with slower disease progression [25]. We previously established a causal link between mHTT expression in the hypothalamus and metabolic and depressive-like phenotypes by the use of adeno-associated viral vectors (AAVs) to selectively overexpress HTT fragments, using groups injected with either a wild-type HTT construct of 18 CAG representing normal HTT (wtHTT 18Q) or a HTT fragment of 79 CAG (mHTT 79Q) [26-28]. Recently, we found that hypothalamus-targeted mHTT 79Q overexpression caused an initial rapid weight gain in the R6/2 mice, but it was insufficient to alter progression to end-stage disease features. In injected R6/2 mice, regardless of whether wtHTT 18Q or mHTT 79Q was overexpressed, the weight loss and loss of key hypothalamic populations were present in R6/2 mice with characteristic disease end-stage phenotype [26]. In addition to prior studies on high caloric diets and the use of transgenic mice to induce weight gain [29-34]; increases in body weight in HD appear to be insufficient to significantly modify disease features in animal models. In line with this, van der Burg and colleagues followed up on results from the original BMI and disease progression study [25] and reported that there is no causal relation between BMI and age of onset in clinical HD [35]. Hence, there is a need for elucidating specific mechanisms underlying HD metabolic changes and how it modifies disease features in HD.

Striatal transcriptional dysregulation is one of the hallmarks of HD [36-38] and specific gene expression changes using qRT-PCR have also been reported in the HD hypothalamus [39-41]. However, large-scale transcriptome analysis has not been performed to identify the hypothalamic alterations after targeted wtHTT and mHTT expression. Therefore, in this study, we used the Affymetrix microarray platform on hypothalamic samples from two different HD mouse models; the transgenic BACHD mice (bacterial artificial chromosome(BAC)-mediated) and the AAV-mediated animal models. BACHD mice have a ubiquitous expression of a full-length mHTT fragment in contrast to the AAV-mediated HD models of WT mice with hypothalamus-targeted overexpression of N-terminal HTT fragments (wtHTT 18Q; AAV-HTT853-18Q or mHTT 79Q; AAV-HTT853-79Q). Both models share the feature of increased food intake and early weight gain but differ in the rate of disease progression and extent of hypothalamic pathology [27, 42]. We analyzed each mouse model separately using age-matched WT littermates as control groups. AAV groups had the highest number of genes in terms of adj.p < 0.05 and range of log2(FC) for limma. Subsequent analysis by Gene Set Enrichment Analysis (GSEA) of the wtHTT 18Q vs. WT and mHTT 79Q vs. WT datasets identified hypothalamic processes that are shared by both datasets, including biosynthesis of sterol in the KEGG-GSEA. Further, while both AAV datasets indicated a mutual impact on the neuroendocrine system, the effect was most pronounced in mHTT 79Q vs. WT; the majority of GO terms were related to neuronal function and specific sets of neurotransmitters, including populations involved in the hypothalamic crosstalk with the periphery. In contrast, GO-GSEA of the wtHTT 18Q vs. WT dataset mainly found processes that could not be interpreted as specifically classified to one specific cell type.

Taken together, our data provide support for transcriptional dysregulation as a significant mechanism of action for mHTT in inducing hypothalamic pathology in HD animal models.

## Results

### Linear Models for Microarray Data (limma) analysis of differential expression in AAV and BACHD datasets

In total, 34760 variables (probe IDs) from the Affymetrix arrays were imported for analysis for each dataset. Limma outputs (Supplemental data 1) are visualized using Volcano plots in Figure 1 with coloring based on a p-value cutoff at 0.05 and abs(log2(FC)) at 1. Limma of the AAV datasets found 4396 (p < 0.05) and 735 (adjusted p (adj.p) < 0.05) variables in 18Q vs WT, 4852 (p < 0.05) and 721 (adj.p < 0.05) in 79Q vs WT, as well as 1475 (p < 0.05) and 0 (adj.p < 0.05) in 79Q vs 18Q. AAV datasets comparing injected mice to uninjected (18Q vs. WT and 79Q vs. WT) showed a skewed distribution with a preference for upregulated genes. Differentially expressed genes were related to the immune system (Figure 1A-B). This feature was not as prominent in the 79Q vs. 18Q dataset that represents a comparison within the vector injected groups (Figure 1C).

**Figure 1.**
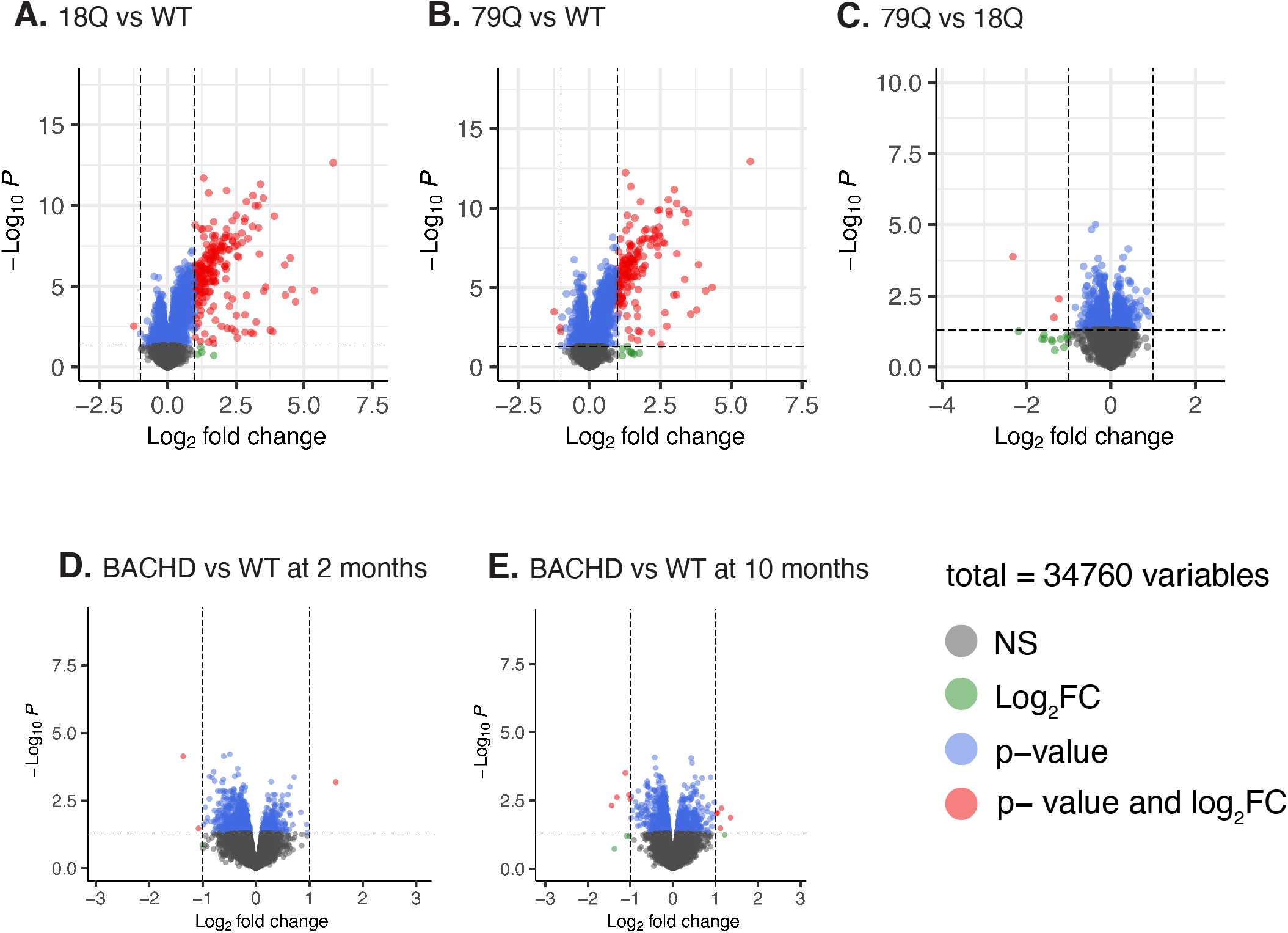
Volcano plots visualizing the p-values as a function of the log2(FC) for the AAV and BACHD microarray datasets compared to age-matched WT controls. Two datasets were analysed for BACHD separated by age to represent early (2 months of age) and late stage of disease (10 months of age) compared with age-matched WT littermates. A total of 34760 variables (probe IDs) from the Affymetrix platform were used as input to the limma analysis. In blue = variables with p-value < 0.05, in gray = non-significant variables (p-value > 0.05 and abs(log2(FC)) < 1) and red = variables with a p-value < 0.05 and abs(log2(FC)) > 1. A log2(fold change) cutoff ± 1 and -log10(p-value) is shown for all graphs. **A)** 18Q vs. WT, **B)** 79Q vs. WT, **C)** 79Q vs. 18Q, **D)** BACHD vs. WT at 2 months of age and **E)** BACHD vs. WT at 10 months of age. 79Q = HTT853-79Q vector, 18Q = HTT853-18Q vector, WT = wild-type, AAV = adeno-associated virus, limma = Linear Models for Microarray Data.

For BACHD datasets, for p < 0.05, 1422 and 1648 probe IDs were identified, respectively, for BACHD 2 months vs. WT and BACHD 10 months vs. WT (Figure 1D-E). Zero variables passed an adj.p < 0.05 adjusting for multiple testing.

### Histidine carboxylase, neuromedin S, and presynaptic dopaminergic marker synaptic vesicle glycoprotein 2c are differentially suppressed after targeted overexpression of mutant HTT

WtHTT and mHTT have been shown in previous studies to exert several different effects in the body (reviewed in [7-9]). Therefore, we further analyzed limma outputs to explore shared and unique effects of HTT on the hypothalamic transcriptome in the AAV-mediated model. First, we excluded probe IDs with “NA” which generated 23456 variables of the total 34760 variables from limma. By sorting for highest log2(FC) for both upregulated (FC > 0) and downregulated (FC < 0) genes, we retrieved the top 20 genes (top 10 upregulated and top 10 downregulated genes). For 79Q vs WT we additionally used an adj.p < 0.05, while no p-value cutoff was used for the 18Q vs WT and 79Q vs 18Q datasets. Heatmaps with hierarchial clustering are shown in Figure 2. For 18Q vs WT, among downregulated in the top 20 list were vasopressin (*Avp*; 18Q vs WT log2(FC) = -0.66, p = 0.09) and oxytocin (*Oxt*; 18Q vs WT log2(FC) = -0.58, p = 0.13) (Figure 2A). *Avp* and *Oxt* were similarly found as p > 0.05 in the 79Q vs WT dataset (*Avp*; 79Q vs WT log2(FC) = -0.56 and p = 0.11, *Oxt;* 79Q vs WT log2(FC) = -0.45 and p = 0.19). Histidine carboxylase (*Hdc*; 79Q vs WT log2(FC*) =* -0.81), neuromedin S (*Nms*; 79Q vs WT log2(FC) = -0.72), stomatin-like protein-3 (*Stoml3;* 79Q vs WT log2(FC) = -0.60) and synaptic vesicle glycoprotein 2c (*Sv2c*) (79Q vs WT log2(FC) = -0.46) were among the top 10 most downregulated (adj.p < 0.05) in the 79Q vs WT dataset (Figure 2B).

**Figure 2.**
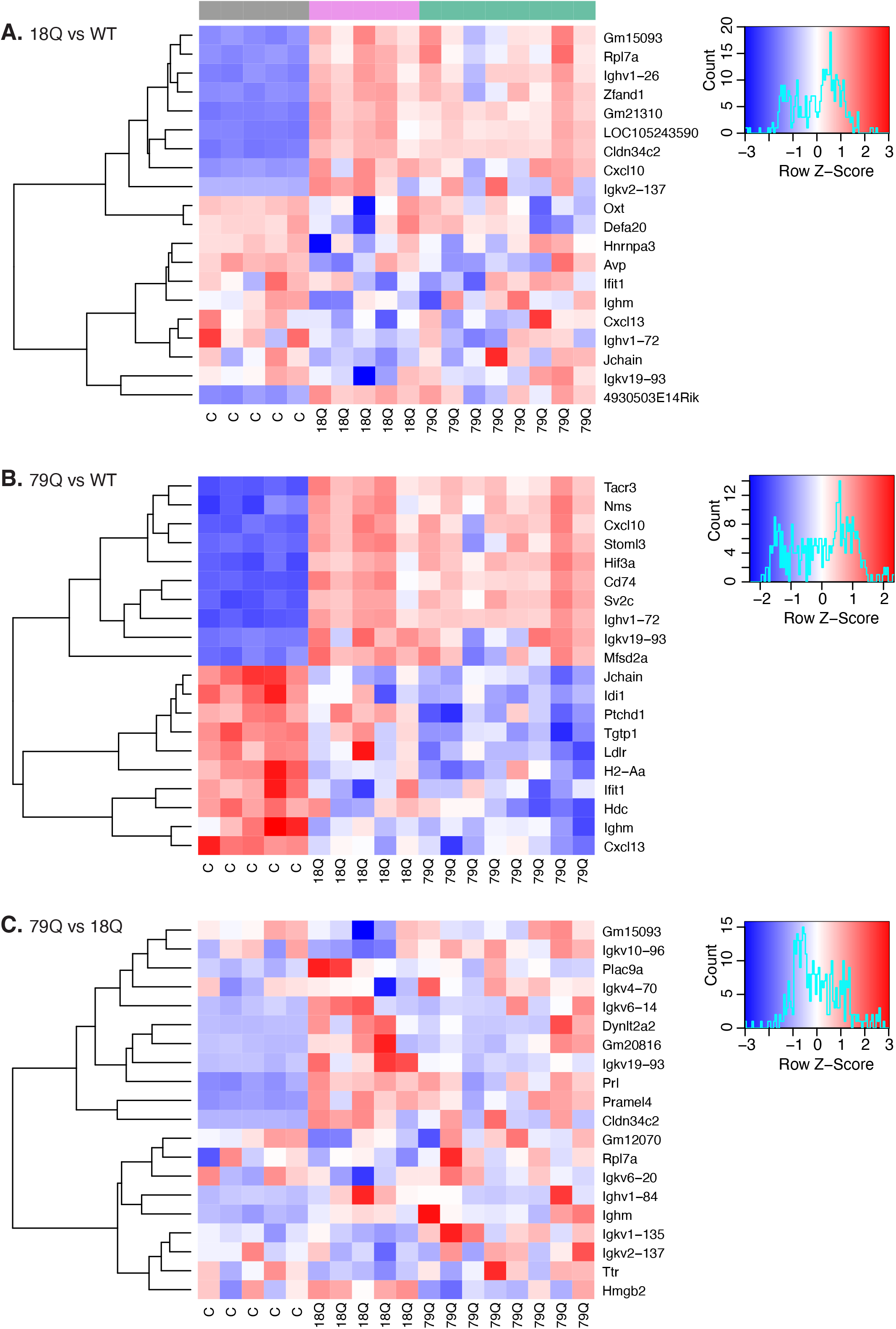
Hierarchial cluster heatmaps visualizing the top-ranked upregulated and downregulated in AAV microarray datasets analyzed by limma. The top 10 upregulated and top 10 downregulated genes are shown for each AAV dataset. For 18Q vs WT and 79Q vs 18Q, no p-value cutoff was used, while for 79Q vs WT all genes displayed are p < 0.05 and adj.p < 0.05. The “C” annotation in the graph corresponds to the WT control group. **A)** Top genes in the 18Q vs. WT dataset, **B)** 79Q vs. WT, and **C)** 79Q vs. 18Q. 79Q = HTT853-79Q vector, 18Q = HTT853-18Q vector, WT = wild-type, AAV = adeno associated virus, limma = Linear Models for Microarray Data.

Comparison of the top upregulated genes ranked by the highest log2(FC) showed that 7/10 were shared between 18Q vs. WT and 79Q vs. WT, all related to the immune system. This was further reflected in the 79Q vs. 18Q dataset (Figure 2C). Among genes were interferon-induced protein with tetratricopeptide repeats 1 (*Ifit1*) an innate immune effector molecule [43], C-X-C motif chemokines (*Cxcl10, Cxcl13*) secreted during inflammatory conditions [44-46] as well as components of antibodies/immunoglobulins (Ig) (*Ighv*; Ig heavy variable, *Igkv*; Ig kappa variable, *Jchain*; joining the chain of multimeric IgA and IgM and *Ighm*; immunoglobulin heavy constant mu) (Supplementary data 1).

### Hypothalamic wild-type and mutant HTT overexpression alter functional neuroendocrine networks and oxidative stress responses

Next, we used GSEA of limma outputs to assess the biological relevance of the altered gene expressions for each comparison. We chose limma-GSEA [47] over single-gene methods as it is advantageous for analyzing and interpreting large-scale datasets. In contrast to single-gene methods with a selective focus on high-scoring genes, GSEA is structured to explore outputs of differential gene analyses (such as limma) in-depth by estimating gene sets and their cross-correlations. In turn, this improves the signal-to-noise ratio. Further, core genes that strongly contribute to enrichments are identified using a subsequent leading-edge analysis [47]. GSEA was performed using R and clusterProfiler (see *Methods*) [48, 49].

First, we performed GSEA of Gene Ontology (GO) for functional annotation where genes are enriched among three classifications; Molecular Function (MF), Cellular Component (CC), and Biological Processes (BP) [48, 49]. For AAV datasets, we found 1306, 1218, and 415 significantly enriched GO terms for 79Q vs. WT, 18Q vs. WT, and 79Q vs. 18Q, respectively (Figure 3A, Supplementary data 2). Of the GO terms shared across all 3 AAV datasets, the majority were related to the immune system; one GO term was related to appetite (“feeding behavior”, GO:0007631) with normalized enrichment score (NES) = -1.90 indicating suppression. The “feeding behavior” leading-edge subset consisted of 42 high scoring genes that formed the core contribution for the enrichment score [47] (leading-edge: tags=32%, list=29%, signal = 28%) (Supplementary data 2).

**Figure 3.**
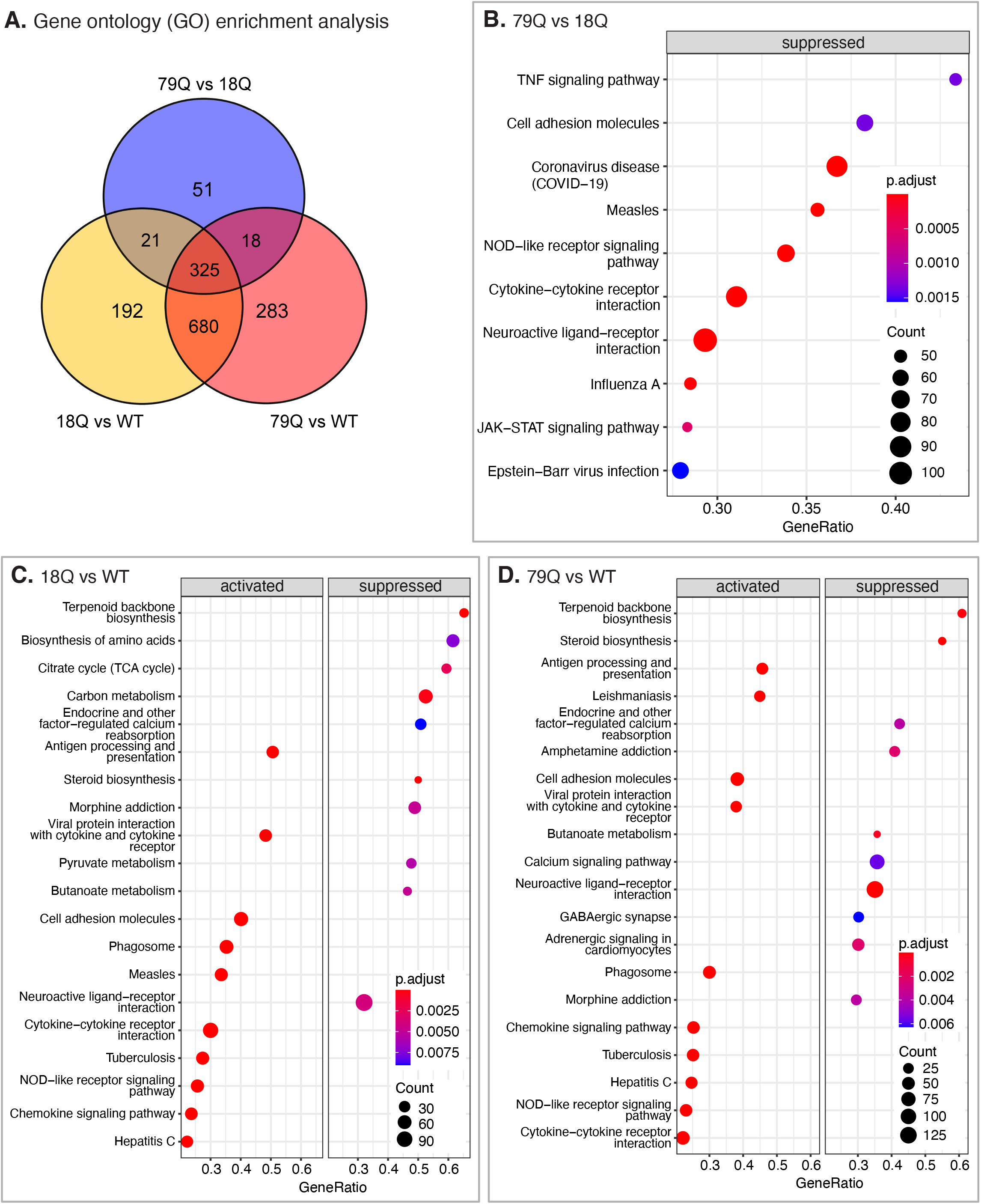
GSEA of gene ontology (GO) and KEGG pathways in AAV groups. **A)** Venn diagram illustrating the relation of GO enrichment between the three AAV datasets. The full list of GSEA-GO outputs can be found in Supplementary data 2. In B-D are the top 10 upregulated and top 10 downregulated/suppressed pathways identified by GSEA of KEGG pathways. **B)** 18Q vs WT, **C)** 79Q vs WT, **D)** 79Q vs 18Q. The full list of GSEA-KEGG outputs can be found in Supplementary data 3. 79Q = HTT853-79Q vector, 18Q = HTT853-18Q vector, WT = wild-type, AAV = adeno associated virus, GSEA = Gene Set Enrichment Analysis, GO = gene ontology

Next, we analyzed unique GO terms in AAV datasets (Figure 3A, Supplemental Table 2). For the vector-injected 79Q vs. 18Q dataset, most GO terms related to processes that could not be specifically attenuated to one specific cell type, and we did not find any immune-system related genes that were unique to only this dataset compared to 18Q vs. WT and 79Q vs WT. For 79Q vs. WT, we found a pronounced set of unique GO terms that could be specifically attenuated to neurons, all NES < 0 that indicate downregulation/suppression. Most neuron-related terms were associated with synaptic transmission, and some annotations defined biological processes involving specific neurotransmitters such as catecholamine, dopamine, neuropeptide Y, and norepinephrine. Further, we found GO terms attenuated to the periphery. First, “gastric acid secretion” (GO:0001696), where we found unique genes constituting the leading-edge subset; neuromedin U (*Nmu*), prostaglandin E receptor 3 subtype EP3 (*Ptger3*), and cholecystokinin receptors A and B (*Cckar, Cckbr*). Second, we found “positive regulation of digestive system process” (GO:0060456), corticotropin-releasing hormone (*Crh*), and tachykinin 1 (*Tacr1*) were among substituents of the leading-edge subset. Opioid receptor-like 1 (*Oprl1*) and *Oxt* were shared among the leading-edge subsets. In contrast, the majority of GO terms in the 18Q vs. WT dataset were not specified by cell type. Instead, we found processes related to ion homeostasis, mitochondria, nitrogen species, and oxidative stress. We further expanded on the 4 GO terms relating to mitochondria that were constituted by large leading-edge subsets (min-max range 31-120 genes), one of the four being “mitochondrial respiratory chain complex assembly” (GO:0033108; leading-edge: 42/81 in subset, tags=52%, list=34%, signal=34%). Among individual genes across all 4 mitochondrial leading-edge subsets (n=189), 65 genes were shared within at least two GO terms (Supplementary data 2).

### Biosynthetic pathways in the hypothalamus are among the topmost affected KEGG pathways in both BACHD and AAV-mediated mouse models

In the 79Q vs. 18Q dataset, we found 38 KEGG pathways using GSEA (Supplemental data 3). The top 20 are shown in Figure 3A. Two pathways in 79Q vs. 18Q that were not found in GSEA of 18Q vs. WT and 79Q vs. WT datasets were “Tyrosine metabolism” (mmu00350) and “Growth hormone synthesis, secretion, and action” (mmu04935), both suppressed (NES < 0). For “Tyrosine metabolism”, the pathway was constituted by 13/37 core genes (leading edge: tags=35%, list=15%, signal=30%), among them monoamide oxidase B (*MaoB*), tyrosinase (*Tyr*), tyrosinase-related protein 1 (Tyrp1), *Ddc, Th* and dopamine beta-hydroxylase (*Dbh*). “Growth hormone synthesis” consisted of 47/114 core genes (leading edge: tags=41%, list=24%, signal=31%). Part of the leading edge subset was growth hormone, its receptor, and releasing hormone (*Gh, Ghr, Ghrh*), insulin-like growth factor 1 (*Igf1*), somatostatin with its receptors (*Sst, Sstr1/2/5*), adenylate cyclases (*Adcy10, Adcy4*), and protein kinase C, alpha (*Prkca*).

For 18Q vs. WT and 79Q vs. WT, GSEA found 102 and 114 KEGG pathways, of which 96 were shared (Supplemental data 3). Several of the shared KEGG pathways were biosynthetic, including “Terpenoid backbone biosynthesis” (mmu00900), “Steroid biosynthesis” (mmu00100), “Butanoate metabolism” (mmu00650), and “Pyruvate metabolism” (mmu00620). The top 20 pathways for 18Q vs. WT and 79Q vs. WT (15/20 shared) are shown in Figure 3B-C.

Expanding KEGG GSEA analysis beyond the top results shown in Figure 3, we next explored the whole KEGG GSEA datasets to find unique pathways for each AAV dataset. When comparing 18Q vs. WT and 79Q vs. WT datasets, none of the unique KEGG pathways could be specifically conferred to the immune system. In 79Q vs. WT (n=18 unique), we instead identified further biosynthetic pathways that were suppressed (NES < 0), such as “Glycerolipid metabolism” (mmu00561), “2-Oxocarboxylic acid metabolism” (mmu01210), and “Ubiquinone and other terpenoid-quinone biosynthesis” (mmu00130), pathways that we did not find in the 18Q vs. WT dataset. In contrast, unique KEGG pathways (n=6) for 18Q vs WT that were not found in the 79Q vs WT GSEA, were activated/upregulated (NES > 0); “Ferroptosis” (mmu04216), “Prolactin signaling pathway” (mmu04917) and “Mitogen-activated protein kinase (MAPK)-signaling pathway” (mmu04010), while “Alanine, aspartate and glutamate metabolism” (mmu00250), “Citrate cycle (TCA cycle)” (mmu00020) and “Propanoate metabolism” (mmu00640) were downregulated/suppressed (Supplemental data 3).

We also performed KEGG GSEA for the BACHD datasets. To note, no significant genes passed adj.p-value < 0.05 in limma when adjusting for multiple testing, however as we showed above for 79Q vs. 18Q with adj.p > 0.05 of all genes, in combination with GSEA, we may still identify pathways that could be considered as biologically relevant, consisting of genes that despite modest expression and lower scores exhibit noteworthy cross-correlation [47]. Results can then further be validated in future studies to verify roles in impacting disease features. There were 3 significantly enriched KEGG pathways for 2 months old BACHD vs. WT datasets (Figure S1A). Two were related to fatty acid (FA) synthesis with NES < 0, indicating suppression of gene sets; “Biosynthesis of unsaturated FA” (mmu01040) and “FA elongation” (mmu00062) with respectively 21/34 core genes (leading-edge: tags = 62%, list=18Q, signal 51%) and 15/29 core genes (leading-edge: tags=52%, list=18%, signal=43%) in the leading-edge subset. The FA-related pathways shared 13 core genes, mainly constituted by acyl-CoA thioesterases (*Acot*), elongation of very long-chain fatty acids proteins (*ELOVL*), and 3-hydroxyacyl-CoA dehydratases (*Hacd*). The third pathway detected in KEGG GSEA was “Glutamatergic synapse” (mmu04724) with NES > 0 and the leading edge subset (25/109) constituted mainly by glutamate receptors, calcium channels, and phospholipases. In 10 months old BACHD vs. WT, there were 13 significant pathways. Immune system-related pathways were prominent. Further, there were pathways related to metabolism and the hypothalamic-pituitary-gonadal axis; “alpha-linolenic acid metabolism” (mmu00592, NES = -1.81) with 9/24 core genes (leading-edge: tags=38%, list=11%, signal=33%) and “Gonadotropin-releasing hormone (GnRH) signaling pathway” (mmu04912, NES -1.79) (leading edge: 12/88 genes, tags=14%, list=4%, signal=13%) (Figure S1B). Phospholipases were part of both leading-edge subsets for both pathways (9/9 and 3/12, respectively) (Supplemental data 3).

### qRT-PCR confirms selective mutant HTT-mediated loss of key enzymes in dopamine and histamine synthesis in the hypothalamus of mice overexpressing mutant HTT

Lastly, using qRT-PCR, we validated a set of candidate genes found in the limma and subsequent GSEA analyses for the AAV groups. As described above, we found significant effects on the neuroendocrine system, biosynthetic pathways including FA and cholesterol, and immune system. Therefore, we focused on key genes and components of each pathway that also are shown in the literature to be altered in HD and/or other neurodegenerative disorders. Expression levels of Vesicular monoamine transporter 2 (VMAT2), a transporter of common neurotransmitters such as dopamine and used as a therapeutic strategy for hyperkinetic disorders, were comparable in 79Q vs. 18Q [50, 51] (Figure 4A). 79Q displayed significant downregulations in hypocretin neuropeptide precursor (*Hcrt*), diazepam-binding inhibitor (*Dbi*), *Ddc, Hdc, and Th* in 79Q compared to 18Q (Figure 4B-D, F-G). Decreased Dbi in cerebrospinal fluid has been reported in HD patients [52]. Dbi plays a role in fatty acyl CoA biosynthesis and long-chain fatty acid metabolism, and lack of Dbi in astrocytes induces hyperphagic obesity due to altered unsaturated FA metabolism [53, 54]. As we throughout the study found multiple immune-system-related factors, therefore, we analyzed Glial fibrillary acidic protein (*Gfap*), an astrocyte marker. However, mRNA levels of *Gfap* were comparable between 79Q and 18Q groups (Figure 4E). *Tacr3* was significantly downregulated in both 79Q and 18Q groups compared to WT controls (Figure 2), and qRT-PCR showed similar expression levels of *Tacr3* (Figure 4H). Other genes included for qRT-PCR, but that we did not find any significant difference in 79Q vs. 18Q were: CREB binding protein (*Cbp*), Inhibitor of DNA binding 4 (*Id4*), Acyl-CoA thioesterases (*Acot8, Acot9*), and Bone morphogenetic protein 4 (*BMP4*) (Figure 4I-M).

**Figure 4.**
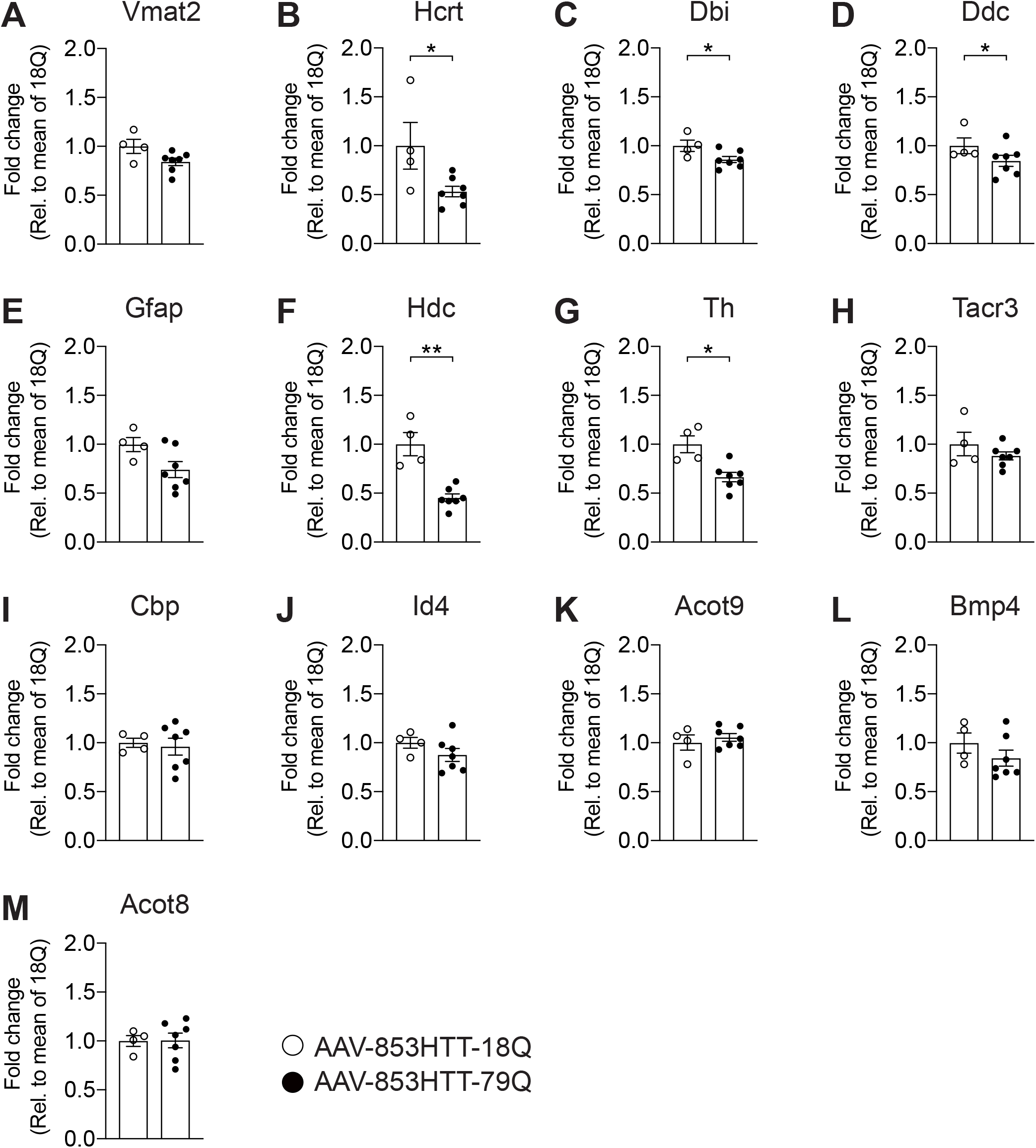
qRT-PCR validation of differentially expressed genes found in AAV groups. Gene expression analysis of **A)** vesicular monoamine transporter 2 (VMAT2), **B)** hypocretin neuropeptide precursor *(Hcrt)*, **C)** Diazepam binding inhibitor (*Dbi*), **D)** dopa decarboxylase (*Ddc*), **E)** glial fibrillary acid protein (GFAP), **F)** histidine decarboxylase (*Hdc*), **G)** tyrosine hydroxylase (*Th*), **H)** tachykinin receptor 3 (*Tacr3*), **I)** CREB-binding protein (*Cbp*), **J)** inhibitor of DNA binding 4 (*Id4*), **K)** acyl-CoA thioesterase 9 (*Acot9*), **L)** bone morphogenetic protein 4 (*Bmp4*) and **M)** acyl-CoA thioesterase 9 (*Acot8*) candidate genes was performed with hypothalamic samples from mice with mutant HTT (79Q) overexpression and wild-type HTT (18Q) overexpression. mRNA expression is presented as relative to the mean of the 18Q group. Non-parametric Mann-Whitney test with *p < 0.05. 79Q = HTT853-79Q vector, 18Q = HTT853-18Q vector, WT = wild-type, AAV = adeno associated virus

We performed qRT-PCR of genes linked to lipid metabolism and GnRH in the BACHD model at 2 months of age, representing the early time point in disease progression. Similar to AAVs, there was no significant difference in mRNA levels of *Acot8* and *Acot9* (Figure 5A, 5H). *Tacr3* was significantly downregulated (Figure 5B). The protein kinase C epsilon (Pkce), phosphorylase kinase regulatory subunit 1 (*Phka1*), fatty acid-2 hydroxylase (*Fa2h*), an inhibitor of DNA binding 2 (ID2), and heat-shock protein 90 (Hsp90) also did not alter in BACHD hypothalamus compared to WT (Figure 5C-G).

**Figure 5.**
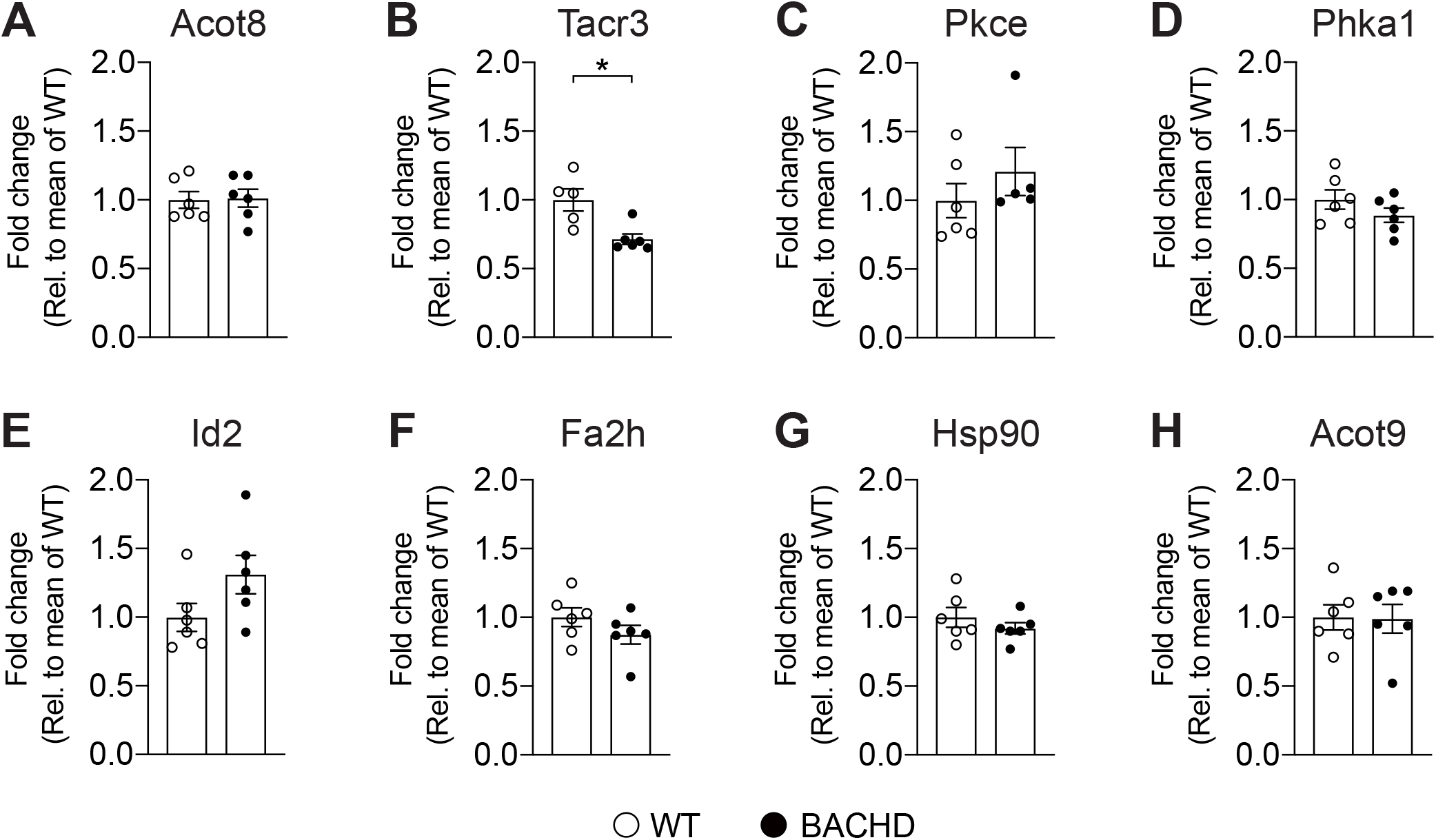
qRT-PCR analysis of candidate genes with roles in lipid metabolism and/or GnRH signaling in 2 months old BACHD. BACHD mice were compared with age-matched WT controls (2 months of age). Data are presented as relative mRNA expression to WT controls. **A)** Acyl-CoA esterase 8 (*Acot*8), **B)** tachykinin receptor 3 (*Tacr3*), **C)** protein kinase C epsilon (*Pkce*), **D)** phosphorylase kinase regulatory subunit alpha 1 (*Phka1*), **E)** inhibitor of DNA binding 2 (*Id2*), **F)** fatty acid 2-hydroxylase (*Fa2h*), **G)** heat shock protein 90 (*Hsp90*) and **H)** Acyl-CoA esterase 9 (*Acot*9). Non-parametric Mann-Whitney test with *p < 0.05. WT = wild-type, GnRH = gonadotropin-releasing hormone

## Discussion

Using Affymetrix microarray profiling for large-scale gene expression analysis, this study aimed to assess changes in hypothalamic transcriptome profiles in mice with overexpression of wtHTT or mHTT in the hypothalamus and the full-length mHTT-expressing transgenic BACHD model of HD. These HD mouse models share a similar metabolic phenotype, namely an increased food intake and body weight [26, 27].

As we in previous studies of the AAV-mediated models were restricted by focusing on individual genes of interest using qRT-PCR, microarray followed by limma-GSEA allowed us to expand on and enhance interpretation of how the hypothalamic gene expression profiles are altered in mice with wtHTT overexpression (18Q) and mice with mHTT (79Q) overexpression. In line with previous work [26, 27, 55], we see that mHTT (79Q) overexpression in the hypothalamus elicits a more pronounced effect on disease mechanisms, such as the local loss of neuropeptides, compared to wtHTT (18Q) overexpression. Among shared top FC-ranked genes for downregulation, *Hdc* encoding the rate-limiting enzyme of L-histidine conversion to histamine was significantly downregulated in 79Q compared to the 18Q group. Histamine is part of the regulatory network of leptin to suppress food intake in the brain [56], and induced *Hdc* deficiency *in vivo* causes increased food intake, susceptibility to high fat diet-induced obesity, and leptin resistance [57-59]. We previously showed that overexpression of mHTT (79Q) in the hypothalamus leads to the development of an obese phenotype with leptin resistance [39]. Therefore, reduced levels of histaminergic input in the hypothalamus could be one of the causative factors for the development of metabolic imbalance that we’ve previously observed in the AAV model [26-28]. GSEA of AAV groups in the present study indicated that this might occur as early as at 4 weeks post-injection. Furthermore, implications for a peripheral link consequently of mHTT overexpression in the hypothalamus were found by GSEA of GO-terms. Peripheral symptoms are present in both clinical HD and animal models (reviewed in [9]). Together with present results, this further verifies the hypothalamus, with its extensive communications in the brain and periphery, as a candidate area to consider in diseases with ubiquitous expression of the mutant protein, as in HD.

Expanding on the feature of mHTT-induced neuropathology and loss of normal HTT function (here modeled by the wtHTT 18Q vector construct), alterations in cholesterol and FA metabolism that provide the main components of the myelin sheath and precursors of steroid hormones [60] are also found in HD [61-63]. Strategies of cholesterol delivery to the HD brain and targeting the key components Sterol-regulatory element-binding protein 2 (SREBP-2) and Sirtuin 2 (SIRT2) in the striatum can ameliorate disease phenotypes [64-66]. Here we found that *Stoml3*, present in cholesterol-rich lipid rafts and associated with dopaminergic transmission [67], was differentially downregulated in 79Q vs. WT and trended towards downregulation in 18Q vs. WT (p < 0.05 but adj p > 0.05). Another suppressed candidate gene in 79Q vs. WT was *Sv2c*, a presynaptic dopaminergic biomarker shown *in vitro* to be upregulated by statins [68-71]. Loss of cholesterol and FA in the hypothalamus would have a critical impact on several biosynthetic processes of the neuroendocrine system, and maybe factors that contribute to the widespread reduction of neuropeptides that we observed in limma-GSEA of the 79Q vs. WT dataset, discussed in the previous section.

Immune-related genes and pathways were prominent in the AAV groups studied here. The extensive use of AAVs for gene therapy and gene transfer strategies [72] is partly due to eliciting minimal inflammation and immunogenicity especially following the virus delivery, compared to other vector types [73-75]. As we in the present study chose to analyze the AAV groups at an earlier time point of phenotype progression (4 weeks post-injection) than prior studies [26, 27], we did expect to see an inflammatory response to the early stage of transgene expression [76]. Therefore, in addition to the 18Q vs. WT and 79Q vs. WT datasets, we included a third dataset assessing specific differences, if any, between the injected mice (79Q vs. 18Q). For the 79Q vs. 18Q dataset, all immune-related terms identified by limma and GSEA were shared with the 79Q vs. WT and 18Q vs. WT datasets. However, exploring the 79Q vs. 18Q dataset in more detail, there were notable differences in FC, including Ig-related components. This would indicate to some degree that the immune-related genes observed in the present study are not only a result of AAV vector expression and tissue damage via transgene delivery into the brain; it may also include diverse effects in response to wtHTT and mHTT overexpression. This has implications for further understanding mechanisms behind metabolic alterations in HD, given the prominent feature of inflammation in obesity and metabolic syndrome [77-79] and prior studies in pre-manifest and manifest HD observing effects of mHTT on inflammatory response and immune cell dysfunction [80-84]. Notably, long-term expression of green fluorescent protein (GFP) using AAVs did not result in any change in body weight in mice [27]. Future studies should be conducted with a focus on more detailed histological and morphological characterization to elaborate on how the local inflammatory environment and its key cell types in the hypothalamus react to increased levels of wtHTT and mHTT.

## Conclusions

Using a microarray platform, we assessed transcriptional alterations in the hypothalamus of two HD mouse models. We have identified novel pathways in the hypothalamus, of which several have also been previously linked to other brain areas in HD. The transcriptional alterations in hypothalamic neuroendocrine neurons were specific to mHTT overexpression in the hypothalamus. In contrast to mHTT, we report that wtHTT overexpression mainly affects processes that can be associated with multiple cell types, of which further studies are needed to elaborate on whether such processes are specific to neurons or also altered in other hypothalamic cell types. Taken together, present results provide implications for validating and designing therapeutic interventions aiming to alter levels of both wtHTT and mHTT in HD. Further studies are warranted to validate the biological roles of processes and pathways reported here on disease features in each HD model.

## Methods

### Ethical considerations

All the mice used in the study were housed in groups and maintained at a 12-hour light/dark cycle with free access to a normal chow diet and water. All the experimental procedures performed in mice were carried out in accordance with the approved guidelines in the ethical permits approved by Lund University Animal Welfare and Ethics committee in the Lund-Malmö region (Ethical permit numbers: 12585/2017, M20-11, M65-13, M135-14).

### Animals

Microarray profiling of the hypothalamic transcriptome was performed in AAV vector-mediated groups of WT mice with targeted expression of wtHTT (AAV-HTT853-18Q) or mHTT (AAV-HTT853-79Q) fragments and BACHD mice that express full-length mHTT (97Q) [42]. Both HD models were compared to age-matched WT controls. All mice used in the study were females from the FVB/N strain.

AAV vector-mediated HD models achieve region-specific overexpression of HTT fragments in the brain through targeted injections using stereotactic surgery. AAV groups were assessed at 4 weeks post-injection; as we showed in our prior studies 4 weeks post-injection was the earliest timepoint for a significant weight gain [26, 27]. The vector constructs used in the present study was a recombinant AAV vector of serotype 5 (rAAV5) carrying an 853 amino acid N-terminal HTT fragment corresponding to either wtHTT (18 CAG repeats; AAV-HTT853-18Q) or mHTT (79 CAG repeats; AAV-HTT853-79Q) under control by the human Synapsin-1 (Syn-1) promoter [85]. Stereotactic injections in the hypothalamus were performed as described previously [27]. In brief, 8 weeks old WT female mice were bilaterally injected, and the surgeries were performed under isoflurane anesthesia. The anterior-posterior (AP), and medial-lateral (ML) stereotaxic coordinates for the hypothalamus were determined according to bregma, and dorsal-ventral (DV) coordinates were calculated from the dura mater. The hypothalamic coordinates were AP = 0.6 mm, ML = 0.6 mm and DV = 5.3 mm. A total viral vector volume of 0.5 μl was delivered in each hemisphere. Following an initial injection of 0.1 μl of viral vector solution, 0.05 μl of viral vectors were delivered in 15 s intervals. Following the injection, the glass capillary was left in the target for an additional 5 min. The vectors and titers were as follows: rAAV5-hSyn-HTT853-18Q: 1.3E+14 GC/ml, and rAAV5-hSyn-HTT853-79Q: 1.2Ex14 GC/ml. A group of WT littermates (uninjected mice) was kept as a control group. Group numbers for each AAV group were as follows: 18Q: n=5, 79Q: n=8, and WT controls: n=5.

BACHD is a transgenic mouse model of HD and ubiquitously expresses a full-length human mHTT (97 CAG repeats; 97Q) [42]. In the study, BACHD mice were assessed at two time points, one group representing the early (2 months of age) and the second group late stages (10 months of age) of the disease in comparison to their WT littermate controls [27, 42]. For 2 months of age, group numbers were the following: BACHD: n=6 and WT: n=6, and for 10 months of age: BACHD: n= 5 and WT: n=3.

### Tissue collection and RNA extraction

Hypothalamic tissue was dissected on ice and snap-frozen in liquid nitrogen after a terminal dose of sodium pentobarbital (600mg/kg, Apoteksbolaget) via intraperitoneal injection. Total RNA was extracted using the RNeasy Lipid Tissue Mini Kit (Qiagen, US) according to the manufacturer’s instructions. RNA concentration and RNA quality measured in terms of RNA integrity number (RIN) were determined using the Agilent 2100 Bioanalyzer (Agilent Technologies, US). Samples with poor RIN (<7) were omitted from further analysis. Microarray analysis was performed on total hypothalamic RNA using the Affymetrix platform (Mouse Gene ST 1.0 array; Thermo Fisher Scientific, US)

### Data pre-processing, limma and gene set enrichment analyses

Analyses were made using R v.4.1.1 [86]. Raw .CEL-files obtained from the microarray analysis using the Affymetrix platform were imported using ReadAffy followed by pre-processing of the raw data set using Robust Multi-Array Averaging (RMA), both functions were from the affy package (v. 1.70.0) [87]. A design matrix was created, illustrating which samples belonged to which analysis group. WT samples were used as a reference level. A linear model was fitted for each gene using lmFit. MakeContrasts was used to specify which groups to compare followed by contrasts.fit to perform the comparison [88]. Subsequently, empirical Bayes smoothing was applied to the standard errors (limma, v.3.48.3) [89]. This resulted in a result matrix for each contrast comparison. ClusterProfiler (v.4.0.5) was used to perform gene set enrichment analysis [48, 49]. GseaGO was used to assess enrichment of gene ontology terms and gseaKEGG to assess enrichment of KEGG pathways. The analysis was performed on the entire gene list obtained from the limma analysis, besides the removal of probe IDs to which no gene could be mapped (“NA”). As organism, mogene10sttranscriptcluster.db was used and mmu for KEGG [90, 91] (KEGG URL: https://www.kegg.jp/ or https://www.genome.jp/kegg/). The reported p-values were further adjusted using the Benjamini-Hochberg procedure to correct for multiple testing. Outputs from all procedures can be found in Supplementary data 1 (limma), Supplementary data 2 (GO) and Supplementary data 3 (KEGG).

### Cluster heatmaps

The limma results files were filtered based on sorting from the highest log2(FC) to the lowest. The top 10 most upregulated and top 10 most downregulated were extracted and plotted in a heatmap using heatmap.2 in gplots (v. 3.1.1) [92]. Coloring was based on each gene’s Z-value.

### qRT-PCR validation of microarray data

To synthesize cDNA, 1µg of RNA from each sample was reverse transcribed using SuperScript IV Reverse Transcriptase (Invitrogen, US) according to the manufacturer’s instructions. Mouse qRT-PCR primers were designed using Primer3Plus software [93]. qRT-PCR reactions were carried out in triplicates following a 3-step amplification protocol using the LightCycler 480 system (Roche Diagnostics, Switzerland). The ΔΔCT method [94] was used to calculate gene expression changes relative to housekeeping genes β-actin and glyceraldehyde 3-phosphate dehydrogenase (GAPDH). Primer sequences are listed in Supplementary Table 1.

### Statistical Analyses

Statistical analysis of qRT-PCR data was performed using Graphpad Prism 7 software (GraphPad Software). Data were analyzed using non-parametric Mann-Whitney U tests with p < 0.05 considered statistically significant.

## Abbreviations

AAV: adeno-associated viral vector
*Avp*: Vasopressin
BAC: bacterial artificial chromosome
BMI: body mass index
*Cckr*: Cholecystokinin receptor
*Cxcl*: C-X-C motif chemokines
*Ddc*: dopa decarboxylase
FA: fatty acid
FC: fold change
GAPDH: glyceraldehyde 3-phosphate dehydrogenase
GFP: green fluorescent protein
*Grh*: Growth hormone
*Hcrt*: orexin/hypocretin protein
HD: Huntington’s disease
*Hdc*: histidine decarboxylase
*HTT*: huntingtin
Ig: Immunoglobulin
*Igf*: Insulin-like growth factor
limma: Linear Models for Microarray Analysis
*Mao*: Monoamide oxidase
NES: normalized enrichment score
*Nm*: Neuromedin
*Oxt*: Oxytocin
*Ptger*: prostaglandin E receptor
qRT-PCR: quantitative real-time polymerase chain reaction
rAAV5: recombinant AAV vector serotype 5
RIN: RNA integrity number
RMA: Robust Multi-Array Average
SREBP: Sterol-regulatory element binding protein
*Stoml3*: Stomatin-like protein 3
*Sst*: Somatostatin
Syn-1: Synapsin-1
*Sv2c*: Synaptic vesicle glycoprotein 2C
*Tacr3*: Tachykinin receptor 3
*Th*: Tyrosine-hydroxylase
*Tyr*: Tyrosinase
WT: wild-type
wtHTT: wild-type huntingtin
mHTT: mutant huntingtin

## Competing financial interests

The authors declare the following competing financial interest(s): SL is currently a full-time employee of Novo Nordisk A/S and holds a minor share portion as part of her employment.

## Author contributions

R.S.K. and Å.P designed the experiments. E.D., A.S.D, N.A., S.L., R.S.K, J.K. performed the experiments. E.D., N.A., A.S.D, and R.S-K, analyzed the data. E.D., R.S.K, and A.S.D wrote the first draft of the manuscript. All authors reviewed the manuscript and approved the final version.

## Acknowledgments

This work was supported by grants from Bagadilico to ÅP; the Swedish Medical Research Council to Å.P. (grant numbers 2010-4500, 2013-3537, 2018/02559); the Province of Skåne State Grants (ALF) to ÅP. ÅP is a Wallenberg Clinical Scholar (Knut and Alice Wallenberg Foundation, #2019.0467). R.S.-K. was supported by The Swedish Society for Medical Research (SSMF) postdoctoral fellowships. We are grateful for the excellent technical assistance provided by Björn Anzelius, Anneli Josefson, Ulla Samuelsson, and Ulrika Sparrhult-Bjork at Lund University.

---

**Supplementary Figure 1.**
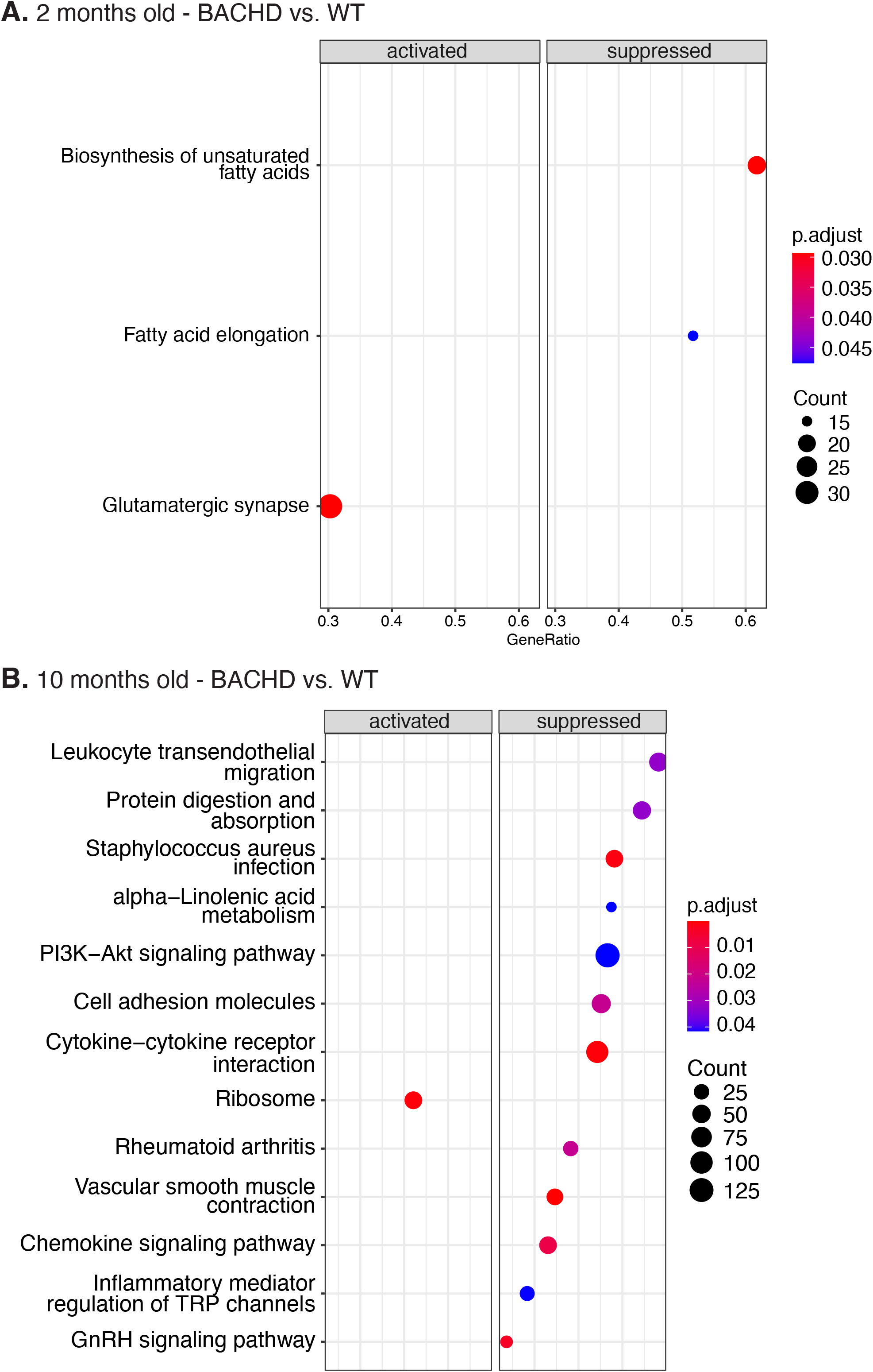

**Supplementary Table 1.**
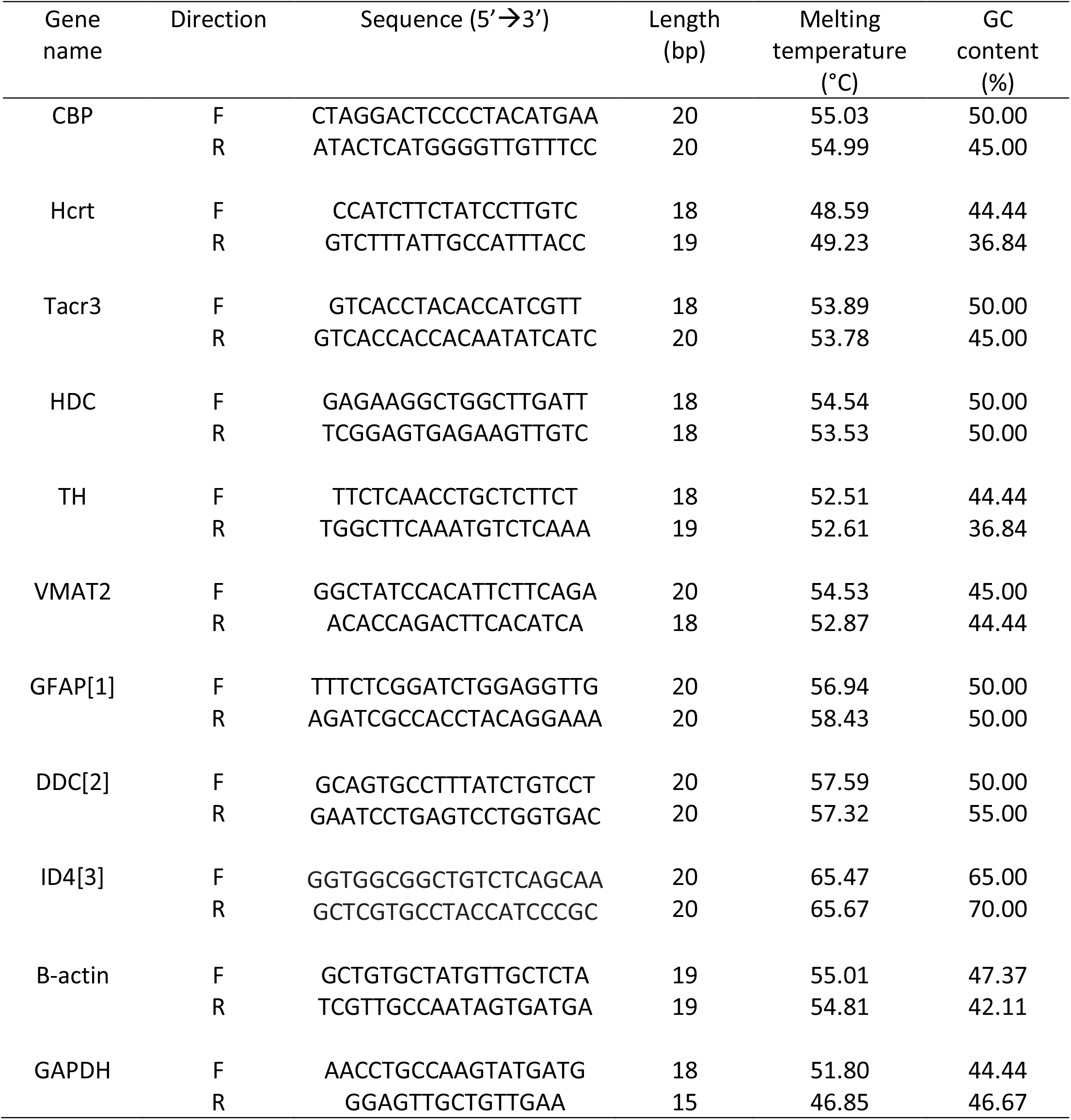

## Notes

### Competing Interest Statement

The authors have declared no competing interest.

